# Leading-edge VASP clusters assemble at sites containing lamellipodin and exhibit size-dependent instability

**DOI:** 10.1101/2020.02.21.960229

**Authors:** Karen W. Cheng, R. Dyche Mullins

**Affiliations:** University of California, San Francisco, CA 94143 USA; Howard Hughes Medical Institute, University of California, CA, 94143 USA

## Abstract

The shape of many eukaryotic cells depends on the actin cytoskeleton; and localized changes in actin assembly dynamics underlie many changes in cell shape. Polymerases of the Ena/VASP family modulate cell shape by locally accelerating actin filament assembly and slowing filament capping. When concentrated into discrete foci at the leading edge, VASP promotes formation of filopodia, but the mechanisms that drive VASP clustering are poorly understood. Here we show that, in migrating B16F1 cells, VASP molecules assemble on pre-existing foci of the adaptor protein, lamellipodin, and that dimerization of lamellipodin is essential for cluster formation. VASP/lamellipodin clusters grow by accumulating monomers and by fusing, but their growth is limited by a previously undescribed, size-dependent instability. Our results demonstrate that assembly and disassembly dynamics of filopodia tip complexes are determined, in part, by a network of multivalent interactions between VASP, lamellipodin, and actin.

## Introduction

Filopodia are thin, finger-like protrusions of the plasma membrane that participate in fundamental cellular processes, including directed migration, substrate adhesion, and cell-cell communication (Blanchoin et al. 2014; Mattila and Lappalainen 2008). Filopodia are defined by morphology rather than function or composition, and growing evidence suggests that different types of filopodia assemble via different mechanisms and employ different sets of actin regulators (Yang and Svitkina 2011; Young et al. 2018; Barzik et al. 2014; Young et al. 2015). One class of filopodia grows from dynamic, lamellipodial actin networks by a process of “convergent elongation”. In this mechanism the growing barbed ends of multiple pre-existing actin filaments converge to a point in the plasma membrane where they are held together by a self-assembling filopodial ‘tip complex’ (Lewis and Bridgman 1992; Svitkina et al. 2003; Mogilner and Rubinstein 2005). The tip complex contains actin polymerases, such as Ena/VASP-family proteins that accelerate filament growth and inhibit filament capping (Hansen and Mullins 2010; Breitsprecher et al. 2008). The spatial constraint imposed on their growing barbed ends causes filaments associated with tip complexes to align into parallel arrays which are crosslinked into tight bundles by the protein fascin (Vignjevic et al. 2003).

Clustering of the polymerase VASP has been identified as a key event in tip complex assembly and an important driver of convergent elongation, but the mechanisms underlying this clustering remain obscure. The Ena/VASP Homology-1 (EVH1) domain of VASP helps determine its subcellular localization by binding proteins that contain an FPPPP (or less commonly LPPPP) motif, such as zyxin, vinculin, Abi1, RIAM, and lamellipodin (Ball et al. 2000; Lafuente et al. 2004; Brindle et al. 1996; Reinhard et al. 1996; Gertler et al. 1996; Prehoda et al. 1999; Krause et al. 2004; Bear et al. 2000; Niebuhr et al. 1997). Using purified proteins and in vitro assays we previously found that lamellipodin, which contains six F/LPPPP motifs and forms membrane-associated dimers (Chang et al. 2013), can co-cluster with VASP on actin filaments (Hansen and Mullins 2015). We also found that clusters of purified lamellipodin recruit VASP and dramatically increase the processivity of its actin polymerase activity. In cells, Lpd recruits VASP tetramers to the leading edge by simultaneously anchoring to the plasma membrane and binding EVH1 domains via multiple FPPPP motifs (Bear et al. 2002; Krause et al. 2004; Hansen and Mullins 2015).

In addition to FPPPP-containing proteins, SH3-containing proteins have also been proposed to play a role in forming functional VASP clusters at the leading edge of migrating cells. In particular, the I-BAR-containing protein, IRSp53, has been shown to recruit purified VASP tetramers to lipid-coated beads *in vitro*, and a Cdc42-activated IRSp53-VASP complex has been proposed as the signaling pathway that clusters VASP in cells (Disanza et al. 2013; Lim et al. 2008). IRSp53 is an attractive candidate for initiating filopodia formation (Welch and Mullins 2002; Ahmed et al. 2010), based on the fact that it can tubulate membranes in vitro and localizes to membrane tubules in live cells (Prévost et al. 2015; Disanza et al. 2013).

At present we know too little about the early stages of VASP cluster formation to know whether lamellipodin or IRSp53 plays a role in the process (Dimchev et al. 2019; Law et al. 2013; Krause et al. 2004; Michael et al. 2010). To address this question, we performed high-resolution time-lapse microscopy of fluorescent actin regulators expressed from their endogenous gene loci in B16F1 mouse melanoma cells. We chose B16F1 cells because they are an excellent model system for studying convergent elongation of leading-edge filopodia. When plated on laminin-coated surfaces, these extend polarized lamellipodia and spread slowly across the surface (Ballestrem et al. 1998). After spreading for 2-3 hours these cells lose their large, polarized lamellipodia and adopt a stable, elongated morphology with abundant focal adhesions and stress fibers. During the initial spreading phase VASP localizes along the dynamic lamellipodial actin network of these cells where it continually coalesces into discrete foci, each of which generates a filopodial actin bundle via the process of convergent elongation (Rottner et al. 1999; Svitkina et al. 2003; Korobova and Svitkina 2008; Mejillano et al. 2004). Our experiments reveal that nascent VASP clusters do not contain IRSp53 but rather arise from pre-existing foci of lamellipodin. By capping actin filaments and over-expressing lamellipodin truncation mutants we demonstrate that VASP clusters are held together by multivalent interactions between lamellipodin, VASP, and free actin barbed ends. Surprisingly, our experiments also revealed that VASP clusters undergo a size-dependent splitting, which appears to set their maximum size. Our data supports a model in which VASP, lamellipodin, and barbed ends make a network of multivalent interactions within the filopodia tip complex to reorganize branched lamellipodial networks into filopodia bundles.

## Results

### Leading-edge dynamics of VASP expressed at endogenous levels in B16F1 melanoma cells

To visualize VASP in live B16F1 cells, we used CRISPR/Cas9 to create a monoclonal cell line expressing VASP-eYFP from the endogenous gene locus (Figure 1A, Supplementary figure 1A). Our knock-in strategy enabled us to study protein localization and dynamics without overexpression artifacts and to quantitatively compare fluorescence intensities between cells and across experiments. Similar to previous studies (Gertler et al. 1996; Reinhard et al. 1992), we found that, during the early stages of cell spreading and migration, VASP-eYFP localized to nascent focal adhesions (Figure 1A, dashed ellipse) as well as the leading edge of advancing lamellipodia, where it often concentrated into discrete foci (Figure 1A, arrowheads; Supplementary figure 1B; Video 1). Compared to B16F1 cells transiently overexpressing GFP-VASP, our engineered cells had significantly fewer discrete foci (Supplemental figure 1C). We occasionally detected the birth of stable VASP foci, which begin as small fluctuations in the leading edge density of VASP (Figure 1B). Consistent with previous studies (Svitkina et al. 2003), we also observed VASP clusters skate laterally along the leading edge, often fusing with each other to form larger clusters (Figure 1B). Stable VASP foci were associated with actin bundles that extend back into the lamellipodial network, and the birth of a VASP focus correlates with the appearance of an associated actin bundle at the leading edge (Figure 1C, Video 2). The fusion of two VASP clusters always resulted in the fusion of their underlying actin bundles via a short-lived forked or lambda-shaped intermediate that rapidly zippers up to form a single, larger bundle (Figure 1C). Overall, the fluorescence intensity of leading-edge actin bundles correlates directly with the intensity of their associated VASP clusters, suggesting that VASP clusters play a role in creating and maintaining their structure (Figure 1D). Some VASP-associated actin bundles form filopodia that protrude several microns beyond the leading edge, while others barely dent the membrane surface. The shorter protrusions are often called ‘microspikes’ to distinguish them from the longer filopodia (Yang and Svitkina 2011), but we observed occasional interconversion between these structures and so, consistent with Svitkina et al. (2003), we refer to them collectively as ‘filopodial actin bundles’.

**Figure 1.**
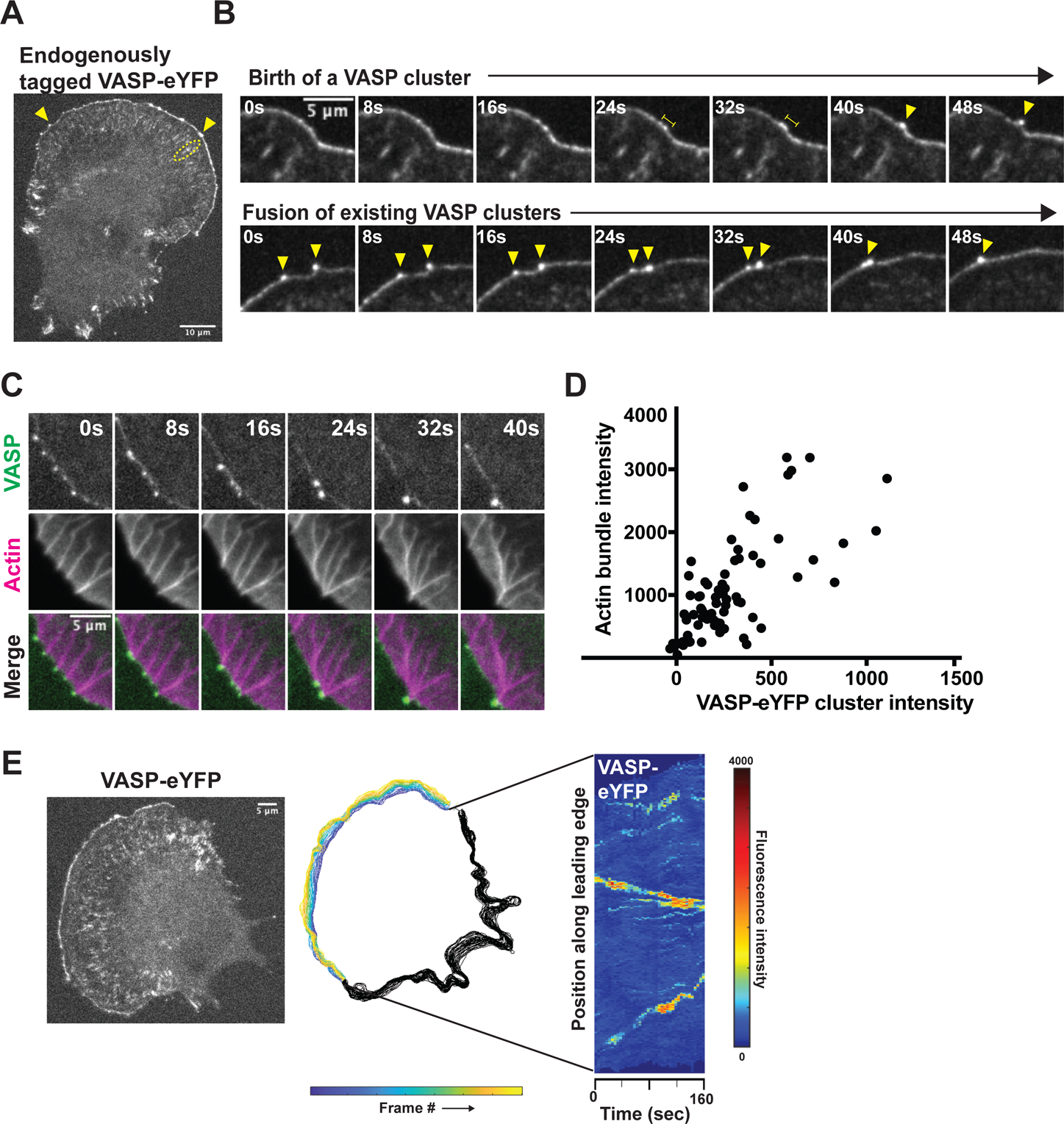
Endogenous VASP-eYFP localization and dynamics during B16F1 cell migration. (A) Representative image of monoclonal, endogenously expressed VASP-eYFP B16F1 cell line generated by CRISPR/Cas9. Yellow arrowheads: VASP clusters; Yellow circle: focal adhesions. (B) Two examples of nascent VASP clusters being born from fluctuations in VASP intensity along the leading edge (top panel) and fusion of existing VASP clusters (bottom panel) by live cell microscopy of the VASP-eYFP B16F1 cell line. Yellow bars: coalescence of VASP-eYFP intensity; yellow arrowheads track VASP-eYFP clusters. (C) Coupled kinetics of VASP-eYFP (green) clustering and SNAP_JF646_-actin bundle (magenta) formation and zippering. (D) Correlation between integrated VASP cluster intensity and underlying actin bundle intensity in 73 foci and 10 cells. (E) Fluorescent image of a representative VASP-eYFP B16F1 cell (left panel) that was used to generate colored outlines of the leading edge position over time (middle panel). Adaptive kymograph map (right panel) re-maps the leading-edge position to the y-axis and time (40 frames or 160 seconds) on the x-axis to visualize VASP-eYFP foci dynamics. Colormap: blue (low fluorescence intensity) to red (high fluorescence intensity).

To more easily track the creation and evolution of VASP-eYFP clusters across the entire leading edge of spreading and crawling cells, we created automated image analysis tools using MATLAB. Briefly, we used VASP-eYFP fluorescence to find the cell edge in every frame of a time-lapse image sequence, and then --based on membrane morphology and dynamics--identified leading-edge lamellipodia for further analysis (Figure 1E, middle panel). For each frame we mapped VASP-eYFP intensity along the current edge onto a vertical line to create a space-time plot, or kymograph (Figure 1E, right panel). Unlike conventional kymographs, which analyze a fixed region of space, these adaptive kymographs follow the advancing and retracting cell edge and simplify analysis of VASP cluster dynamics by: (1) removing membrane movement; (2) reducing the number of spatial dimensions; and (3) mapping time onto space, enabling us to rapidly identify key events in the life cycle of VASP clusters across many cells. Key events include the birth of nascent clusters; lateral skating of mature clusters; and fusion of colliding clusters.

### Lamellipodin is a stoichiometric component of leading-edge VASP foci

The earliest observable event in filopodial bundle formation is the clustering of VASP molecules into discrete foci, so we aimed to identify additional molecules involved in VASP clustering. Multivalent EVH1-binding proteins Mig10, RIAM and lamellipodin (collectively, the MRL family of proteins) associate with VASP in leading-edge lamellipodia, but previous studies differ regarding the association of MRL proteins with VASP foci. We, therefore, asked whether EVH1-binding proteins might also localize to VASP clusters at the leading edge. Krause et al. (2004) initially observed lamellipodin at the tips of filopodial protrusions and co-localized with Mena and VASP foci at the leading edge, while Disanza et al. (2013) claimed that MRL proteins do not localize into discrete foci at filopodia initiation sites.

To investigate the involvement of MRL-family proteins with VASP foci at the leading edge, we created a double CRISPR knock-in B16F1 cell line expressing both VASP-eYFP and a fluorescent lamellipodin fusion protein (Lpd-tdTomato) from their endogenous loci (Figure 2A). Consistent with the original report of Krause et al. (2004), we found that Lpd-tdTomato localizes to lamellipodia and concentrates in VASP-rich foci (Figure 2A, arrowheads). Remarkably, the ratio of time-averaged, integrated intensities of VASP and Lpd within leading-edge foci was approximately constant (R^2^= 0.90) across 58 stable foci observed in four independent experiments using the same imaging conditions (Figure 2C, black circles). In contrast, the fluorescence intensities of Lpd and VASP measured along the entire leading edge were much less correlated (Supplemental figure 2A). These results reveal that, even though lamellipodin and VASP are not present at a constant ratio along the entire leading edge, they are present at a more or less fixed stoichiometry in stable VASP foci.

**Figure 2.**
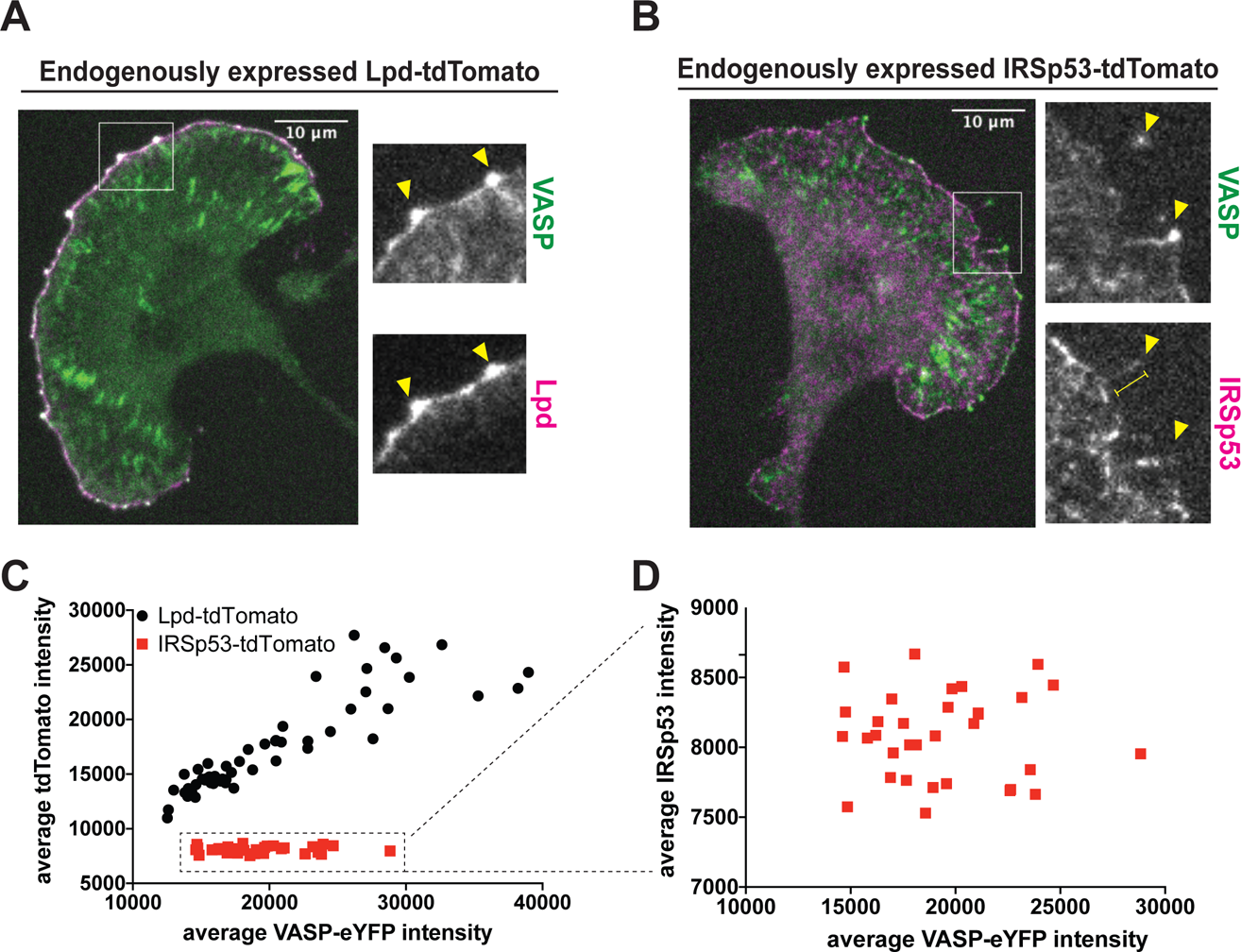
Differential localization of the VASP binding partners, lamellipodin (Lpd) and IRSp53, at the leading edge. (A) Representative image of monoclonal, endogenously expressed VASP-eYFP and Lpd-tdTomato B16F1 cell line shows strong colocalization of the two proteins at VASP foci. Arrowheads: VASP-eYFP foci. (B) Representative image of monoclonal, endogenously expressed VASP-eYFP and Lpd-IRSp53 B16F1 cell line shows poor colocalization of the two proteins at VASP foci. Arrowheads: VASP-eYFP foci; yellow bar: filopodia shaft. (C) The time-averaged integrated intensities of VASP-eYFP and Lpd-tdTomato within leading edge foci are tightly correlated (black circles). A plot of average Lpd intensity versus VASP intensity fits a straight line (R^2^=0.90). In contrast, the integrated intensities of VASP-eYFP and IRSp53-tdTomato within leading edge foci are not correlated (red squares; R^2^=0.00014) (D). Zoomed plot of the VASP-eYFP and IRSp53-tdTomato dataset shows the lack of correlation in their intensities.

### IRSp53 does not persistently localize to leading-edge VASP foci

Disanza et al. (2013) proposed that IRSp53, an I-BAR domain-containing protein found at the plasma membrane, directly interacts with VASP tetramers and induces them to form stable clusters. Other studies, however, suggest that the connection between VASP and IRSp53 may be less direct (Sudhaharan et al. 2019; Nakagawa et al. 2003), and so to determine whether IRSp53 is present in the early stages of VASP cluster formation, we used CRISPR/Cas9 to create a double knock-in B16F1 cell line, expressing both IRSp53-tdTomato and VASP-eYFP from their endogenous loci (Figure 2B). We first used an antibody against IRSp53 to confirm that our IRSp53-tdTomato displays the same localization as untagged endogenous protein (Supplemental figure 3A). In both control and IRSp53-tdTomato-expressing cells, the anti-IRSp53 antibody recognized discrete patches distributed along the plasma membrane and concentrated in the shafts of filopodia, but did not obviously co-localize with clusters of VASP-eYFP (Supplementary figure 3A, B). Similarly, we observed IRSp53-tdTomato fluorescence enriched in patches along the plasma membrane, and often concentrated in the shafts of mature filopodia (Figure 2B, arrowheads). Unlike lamellipodin, however, IRSp53-tdTomato was not a stable component of VASP foci, and the correlation between the time-averaged, integrated intensities of VASP and IRSp53 in leading-edge foci (R^2^= 0.00014) was not significant (Figure 2C, D; red squares). Finally, in addition to tagging IRSp53 at the endogenous gene locus, we also transiently over-expressed IRSp53 fused at its N-terminus to mRuby2. Over-expression of this fluorescent fusion protein induced formation of abundant filopodial protrusions containing mRuby2-IRSp53, which localized along their shafts but rarely extended out to the filopodia tips (Supplemental figure 3C, arrowhead). Only a few of these ectopic, IRSp53-rich filopodia contained detectable VASP-eYFP at their tips. The majority of VASP-eYFP in these cells remained associated with the dynamic lamellipodial actin network, where it formed stable foci that contained no detectable fluorescent IRSp53. Together, these localization data indicate that IRSp53 is not a stable component of leading edge VASP foci.

**Figure 3.**
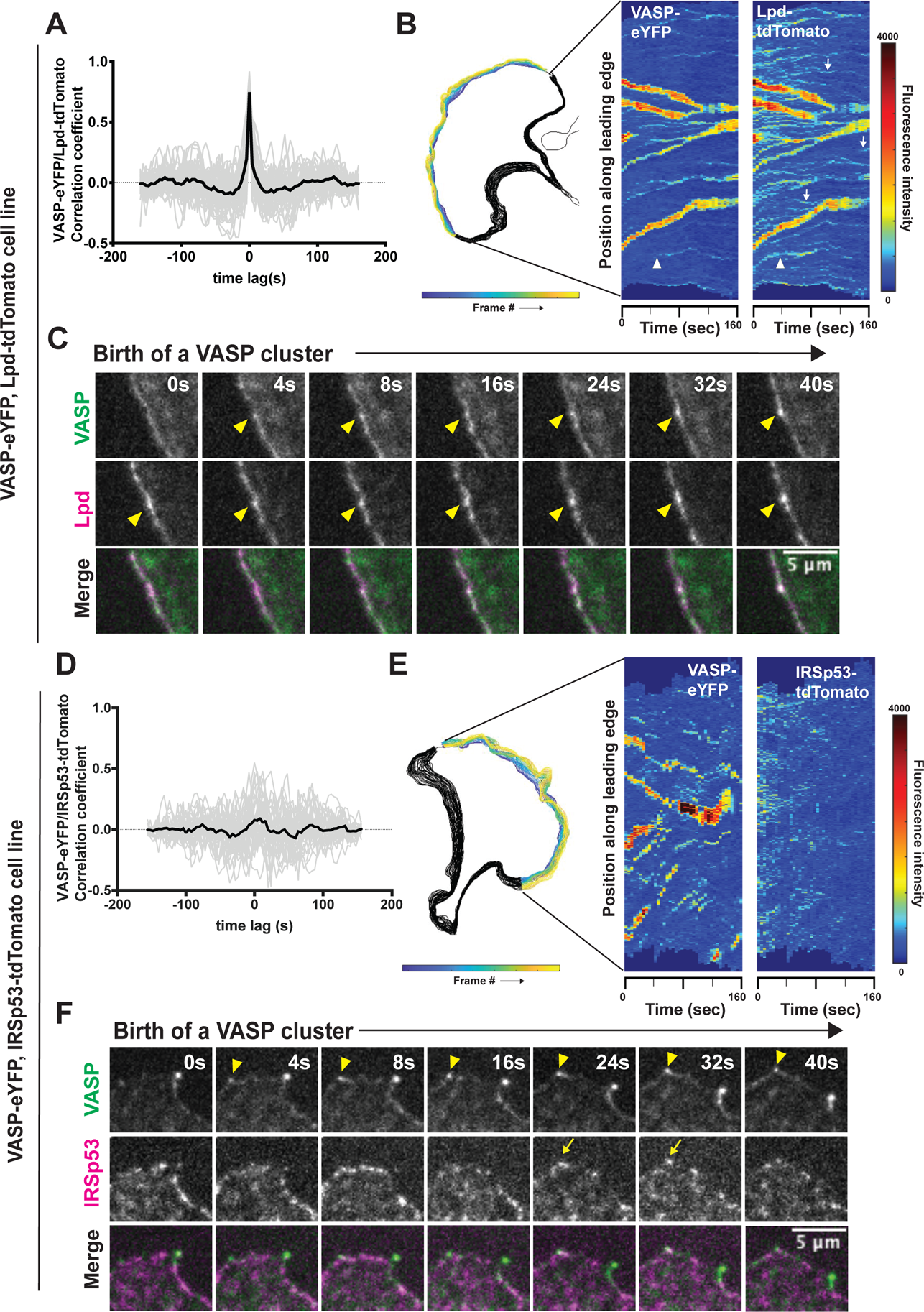
Dynamics of VASP clustering with its leading-edge binding partners, Lpd and IRSp53. (A) Temporal cross-correlation analysis of fluctuations in VASP and Lpd intensity at leading-edge foci. Gray lines are individual cross correlation measurements and the solid black line, which is symmetrical about zero, is the average of individual traces from 46 different clusters in 10 cells. Peak cross correlation occurs at time lag = 0s with a correlation coefficient of r = 0.75. (B) Cell outline of changing leading edge positions over 40 frames (left). Adaptive kymograph map (right) displaying VASP-eYFP and Lpd-tdTomato dynamics over time (40 frames; 160 seconds). (C) Frames from time-lapse microscopy of VASP clusters being born from small pre-existing Lpd clusters (arrowheads) at the leading edge in double knock-in VASP-eYFP(green)/Lpd-tdTomato (magenta) B16F1 cells. (D) Temporal cross correlation of VASP-eYFP and IRSp53-tdTomato fluorescence signal from 34 clusters in 7 cells. Peak cross correlation occurs at time lag = 8s with a correlation coefficient of 0.09. (E) Outline of changing leading edge positions over 40 frames (left). Adaptive kymograph map (right) displaying VASP-eYFP foci and IRSp53-tdTomato dynamics over time (40 frames; 160 seconds). Colormap: blue (low fluorescence intensity) to red (high fluorescence intensity). (F) Frames from time-lapse microscopy of VASP clusters (arrowhead) forming with no detectable accumulations of IRSp53-tdTomato until after the VASP foci is formed (arrow).

### Dynamics of Lpd and IRSp53 within leading-edge VASP foci

We next compared the leading-edge dynamics of VASP with those of its interaction partners, Lpd and IRSp53. For this analysis, we computed the cross-correlation between fluorescence intensities of VASP-eYFP and Lpd-tdTomato (or IRSp53-tdTomato) within individual VASP clusters over a time window of 160 seconds. Cross-correlation of VASP and Lpd fluorescence intensities revealed that, within leading-edge foci, temporal fluctuations of the two proteins are strongly coupled. The peak correlation between the two molecules (r = 0.75) occurs at a time lag of 0 seconds, implying that (within the temporal resolution of our time-lapse sequences) fluctuations in one molecule do not systematically lead or lag fluctuations in the other (Figure 3A). From the adaptive kymographs we identified numerous discrete foci of lamellipodin that lack a corresponding accumulation of VASP-eYFP (Figure 3B, white arrows). These Lpd-only foci were generally smaller, more transient, and exhibited less lateral (skating) movement than foci that contain both VASP and lamellipodin (Figure 3B, Supplemental Figure 2B). Further examination revealed that nascent VASP foci emerge from these small, pre-existing Lpd-only foci (Figure 3B, white arrowhead). We corroborated this observation with the time-lapse movies of the corresponding VASP-lamellipodin clusters, which clearly show that an increased local density of lamellipodin precedes the concentration of VASP into a detectable focus (Figure 3C, arrowhead).

Next, to detect potential transient interactions with IRSp53, we also examined whether fluctuations of VASP and IRSp53 intensities were temporally correlated. The maximum cross-correlation occurred with a time delay of 8 seconds which surprisingly suggests that VASP intensity may weakly drive fluctuations in IRSp53 intensity and not the other way around, though the correlation coefficient (r = 0.093) was quite low (Figure 3D). Importantly, this analysis establishes that IRSp53 accumulation at the leading edge is unlikely to robustly drive VASP accumulations resulting in foci formation. Furthermore, comparison of adaptive kymographs displaying VASP-eYFP and IRSp53-tdTomato dynamics along the leading edge of spreading cells revealed no stable spatial co-localization of the two proteins (Figure 3E). We also could not detect IRSp53 persistently concentrated in either nascent or stable VASP clusters at the leading edge (Figure 3F). In fact, consistent with our temporal analysis, we only occasionally detected IRSp53 concentration in VASP foci after the VASP focus was already formed (Figure 3F, arrow at 24 seconds), suggesting that the VASP focus was recruiting IRSp53 to sites of filopodia initiation and that IRSp53 is not required for the formation of VASP clusters.

### VASP/Lpd clusters undergo size-dependent splitting events

Similar to previous studies of VASP clusters and leading edge microspikes (Svitkina et al. 2003; Oldenbourg et al. 2000), we observed that VASP-lamellipodin foci grow by both continuous incorporation of molecules along the leading edge and by fusion with pre-existing foci. Because these foci remain dynamic and dispersed along the leading edge rather than accumulating into a single large blob, we hypothesized that, in addition to steady growth and occasional fusion, the foci must also undergo shrinkage and/or fission. Consistent with this idea, we identified abrupt and simultaneous drops of both VASP and Lpd intensity within individual foci (Figure 4C, red arrows). These sudden drops in fluorescence intensity turned out to reflect splitting events in which a large bolus of VASP and Lpd abruptly detaches from the original VASP/Lpd cluster (Figure 4A, yellow arrowhead; Video 5). Rather than splitting laterally along the leading edge, however, the direction of fission was approximately orthogonal to the leading edge, with the detaching piece of VASP and Lpd moving inward toward the cell body. As splitting progresses, the inward-moving cluster of molecules begins to lose Lpd, eventually becoming a diffuse mass of VASP alone (Figure 4A, 32 seconds). The portion of the original VASP/Lpd cluster that remains associated with the leading edge exhibited the same dynamics and VASP/Lpd ratio as before the fission event, emphasizing the robustness of the mechanism that maintains the stoichiometry between VASP and lamellipodin in the foci (Figure 4A-C; see Supplemental figure 4A for more examples). Additional kymograph analysis of splitting events revealed that the detached piece of VASP does not treadmill backward with actin retrograde flow, but remains stationary in the laboratory frame of reference as the leading edge moves forward (Figure 4B). In some examples, VASP shed from a leading edge focus adopted the elongated shape of a nascent focal adhesion, and remained in the same spot for the duration of the movie (Figure 4A, arrowhead at 32 seconds, Figure 4B).

**Figure 4.**
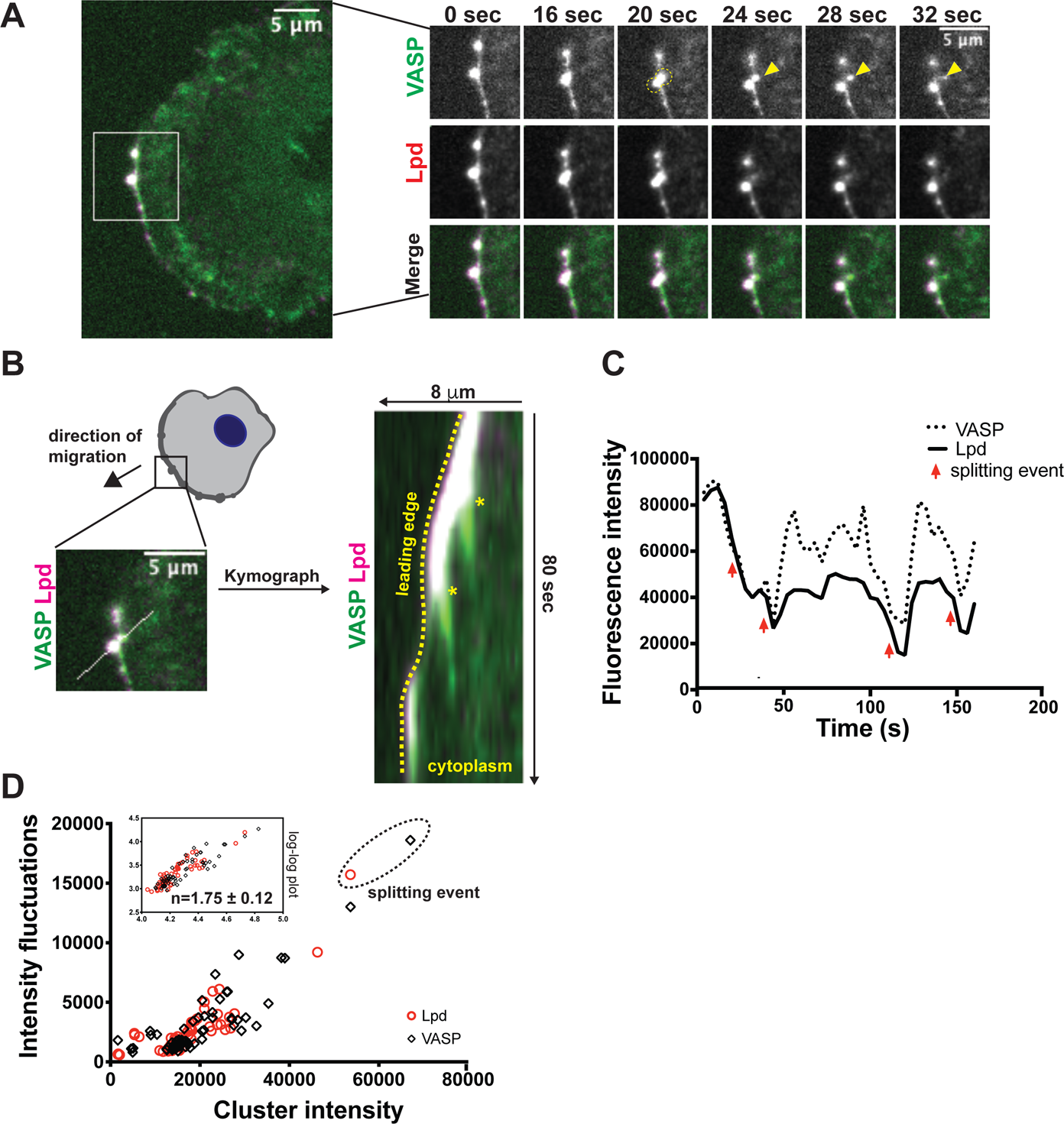
VASP/Lpd clusters display size-dependent splitting behaviors. (A) Image (left panel) of double knock-in VASP-eYFP(green)/Lpd-tdTomato(magenta) B16F1 cell with ROI indicating the VASP/Lpd cluster that undergoes a splitting event in which VASP material splits from the parental cluster (right panels, yellow arrow). (B) Kymograph of a ROI bisecting both the main VASP/Lpd cluster and secondary VASP clump that is shed indicates that the shed VASP does not move relative to the lab frame of view (asterisks). Dotted yellow line outlines the leading edge. (C) Intensity plot profile of the fluorescence intensity of a single VASP/Lpd focus from Figure 4a. Red arrows indicate simultaneous drops in both VASP and Lpd fluorescence intensity which correlate to splitting events. (D) Fluctuations in VASP and Lpd were quantified by plotting the standard deviation in fluorescence intensity of VASP (black diamond) and Lpd (red circle) foci versus its time-averaged VASP (or Lpd) intensity. Inset: log plot of standard deviation vs. time-averaged VASP/Lpd intensity fits a line with slope = 1.75±0.12.

To further explore cluster disassembly, we quantified fluctuations in the intensity of VASP/Lpd clusters over time. By plotting the intensity fluctuations as a function of cluster size, we instead found that VASP/Lpd clusters become more *unstable* as they grow (Figure 4D). In fact, across all of our experiments, size fluctuations are proportional to 1.75-power of the average cluster size (1.75±0.12; Figure 4D, inset: log-log plot). This size-dependent instability effectively limits the maximum size to which VASP/Lpd clusters can grow. Furthermore, we linked the size-dependent instability of VASP/Lpd clusters with the dramatic splitting events that we described above (Figure 4D, circled), as the extremely large clusters in the size distribution plot appear to shrink through the splitting events we observed in cells. To verify that this inverse relationship between cluster size and stability is independent of the specific analysis method, we analyzed the data using an alternative strategy. For each cluster, we plotted the cluster size at each time point (40 time points total) as the vertical scatter (black circles) and as a function of its corresponding time-averaged cluster size (Supplemental figure 4A). Consistent with the original fluctuation analysis method, the vertical scatter (representing increased intensity fluctuations for each VASP/Lpd cluster increased as the cluster size increased. In total, this data reveals that VASP/Lpd clusters become more unstable as they grow in size, which is a novel mechanism for VASP/Lpd cluster size control at the leading edge.

### Lpd requires membrane-targeting and multiple EVH1-binding sequences to incorporate into VASP clusters

To determine which regions of lamellipodin mediate interaction with leading-edge VASP clusters, we transiently overexpressed various fluorescently labeled (SNAP-tag with JF646 ligand) Lpd mutants in our double knock-in B16F1 cells (Figure 5A). First, we overexpressed a N-terminal fragment of Lpd (1-529aa; ‘N-term only’) comprising a coiled-coil motif, Ras-association (RA), and plekstrin homology (PH) domains, which mediate dimerization, G-protein binding, and membrane-localization respectively. This construct lacks VASP binding sites, displays typical B16F1 plasma membrane localization, and fails to incorporate into VASP/Lpd clusters (Figure 5B control vs Figure 5G). Similarly, a C-terminal truncation of Lpd (850-1250aa; ‘C-term only’) --which contains all of the known VASP and actin binding sites (Hansen and Mullins 2015), but lacks G-protein-binding, membrane-targeting, and dimerization sequences--also fails to incorporate into VASP clusters. Perhaps not surprisingly, the localization of C-term Lpd was mainly cytoplasmic, and the protein did not concentrate significantly on leading edge membranes like the endogenously tagged, full-length Lpd-tdTomato (Figure 5H). To determine the role of membrane binding in cluster formation, we fused an unrelated membrane-targeting sequence (the N-myristoyl transferase target sequence GCIKSKGKDSA from the Src-family kinase Lyn (Inoue et al. 2005) to the C-term of Lpd to make Lyn_11_ C-term Lpd. Although Lyn_11_ C-term Lpd now localized tightly to the leading edge, it also failed to incorporate into VASP/Lpd clusters (Figure 5D). Furthermore, we found that, not only does it fail to incorporate into clusters, it effectively inhibited formation of clusters by endogenous VASP and lamellipodin at the leading edge (Figure 5D, Figure 5E vs Figure 5C). This dominant-negative effect of Lyn11 C-term Lpd required its ability to bind VASP, as the VASP binding mutant (F/LPPPP_6_→ APPPP_6_), which does not bind VASP has no effect on VASP/Lpd cluster formation (Figure 5I).

**Figure 5.**
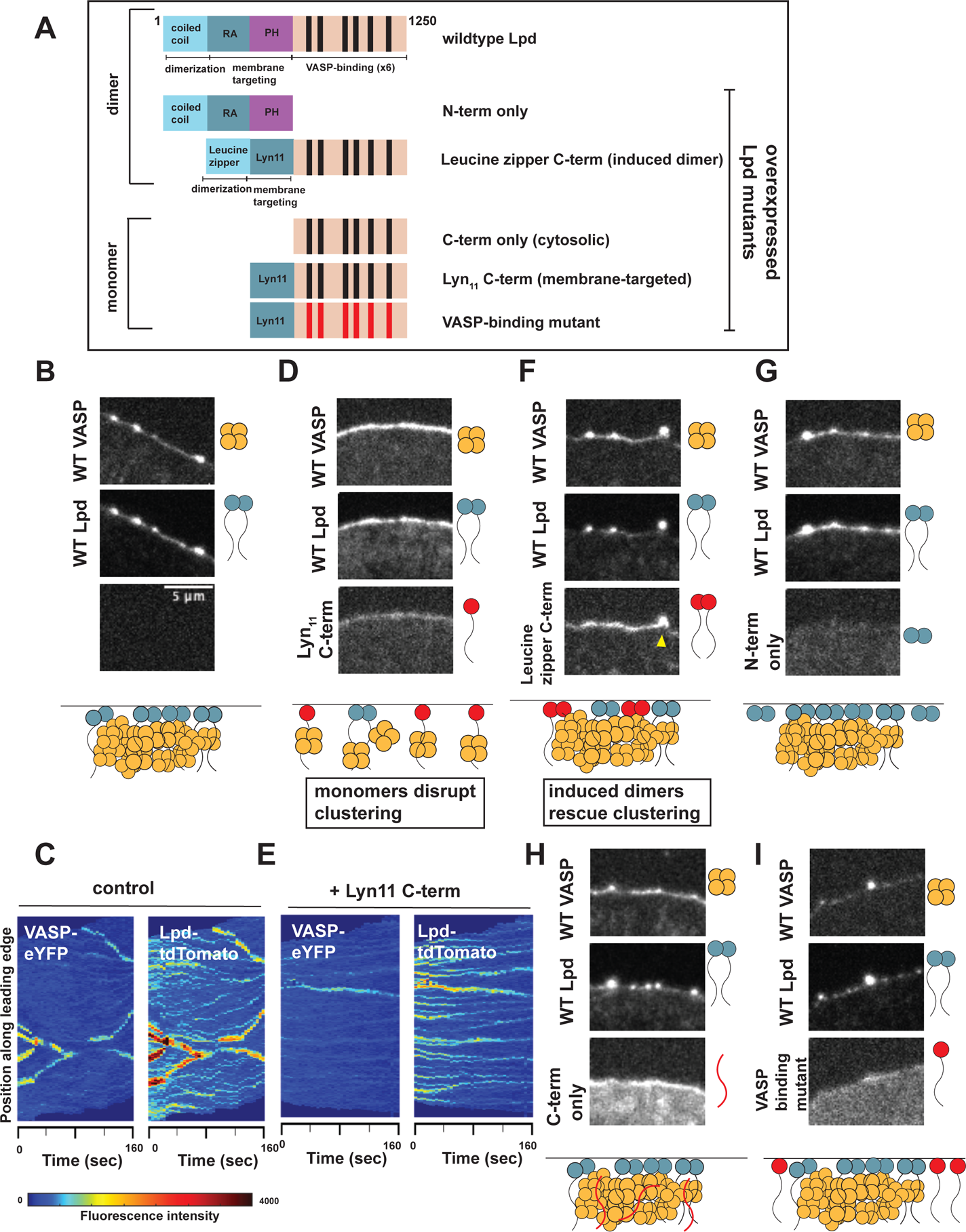
Lpd incorporation into dynamic VASP foci depends on its number of VASP binding sites. (A) Schematic of the domain architecture of wildtype and various Lpd mutants that were over-expressed in B16F1 cells already endogenously expressing VASP-eYFP and Lpd-tdTomato. Leucine zipper (LZ) was used to orthogonally induce dimerization of the Lpd C-term; Lyn11 peptide motif was used to replace the N-term and orthogonally target C-term Lpd to the plasma membrane; VASP binding mutant: F/LPPPP_6_→ APPPP_6_. (B-I) Representative images from three-color microscopy of endogenous VASP-eYFP (top panel) and Lpd-tdTomato (middle panel) and various overexpressed Lpd mutants (bottom panel). All Lpd mutants were over-expressed using a CMV promoter to drive expression of the N-terminal SNAP-Lpd (with Janelia Fluor 646 SNAP ligand) fusion protein. Cartoon schematic below describes the effect that overexpression of each SNAP-Lpd mutant (red) has on endogenous VASP(orange)/Lpd(blue) clustering. Yellow arrow: example in which the leucine zipper C-term construct is able to incorporate into VASP/Lpd clusters. Adaptive kymographs display leading edge VASP/Lpd clusters in (C) wild type double knock-in cells and (E) double knock-in cells with overexpressed monomeric Lyn11 C-term, which has a dominant negative effect.

Finally, we tested the role of dimerization in Lpd clustering by engineering a homo-dimeric leucine-zipper into the C-terminal lamellipodin truncation mutant, to form an induced dimer -- ‘leucine zipper C-term’. The combination of leucine zipper-mediated dimerization and myristoylation-driven membrane localization now partially rescued the ability of the C-terminal tail of Lpd to incorporate into endogenous VASP/Lpd clusters (Figure 5F, yellow arrowhead). Consistent with the finding that only dimerized Lpd incorporates into leading-edge VASP-lamellipodin clusters, we observed that the induced dimer rescued clustering of endogenous VASP and lamellipodin. The ability of the membrane tethered, monomeric Lpd construct to disrupt VASP-lamellipodin clusters argues that these clusters are held together, in part, by multivalent EVH1-FPPPP interactions. Collectively, these data indicate that Lpd incorporation into VASP clusters requires membrane targeting, dimerization, and multiple VASP-binding sites, but does not require interaction with specific small G-proteins or phospholipids.

### Actin filaments with free barbed ends are required to form and maintain VASP/Lpd clusters

Because VASP associates with barbed ends of growing actin filaments at the leading edge, we wondered whether free barbed ends were also required to maintain stable VASP/Lpd clusters. We, therefore, treated B16F1 cells expressing VASP-eYFP and Lpd-tdTomato with Cytochalasin D (CD), a small molecule that rapidly caps free barbed ends. Within 100 seconds of drug treatment, VASP-lamellipodin clusters fell apart (Figure 6A, Video 6). Cluster dissolution was dramatic and proceeded in two steps. First, VASP translocated inward, away from the leading edge, leaving small lamellipodin clusters intact at the leading edge (Figure 6A, B). Next, the lamellipodin-only clusters remaining on the membrane slowly dissolved and lamellipodin redistributed along the membrane (Figure 6A, 200 seconds). The relative change in clustering was quantified as the standard deviation normalized to the mean fluorescence intensity of the cell (Figure 6C, D). The redistribution of Lpd into an even line along the membrane in the absence of VASP and barbed ends suggests that Lpd requires the presence of VASP and barbed ends at the membrane to be held together in a higher-order structure.

**Figure 6.**
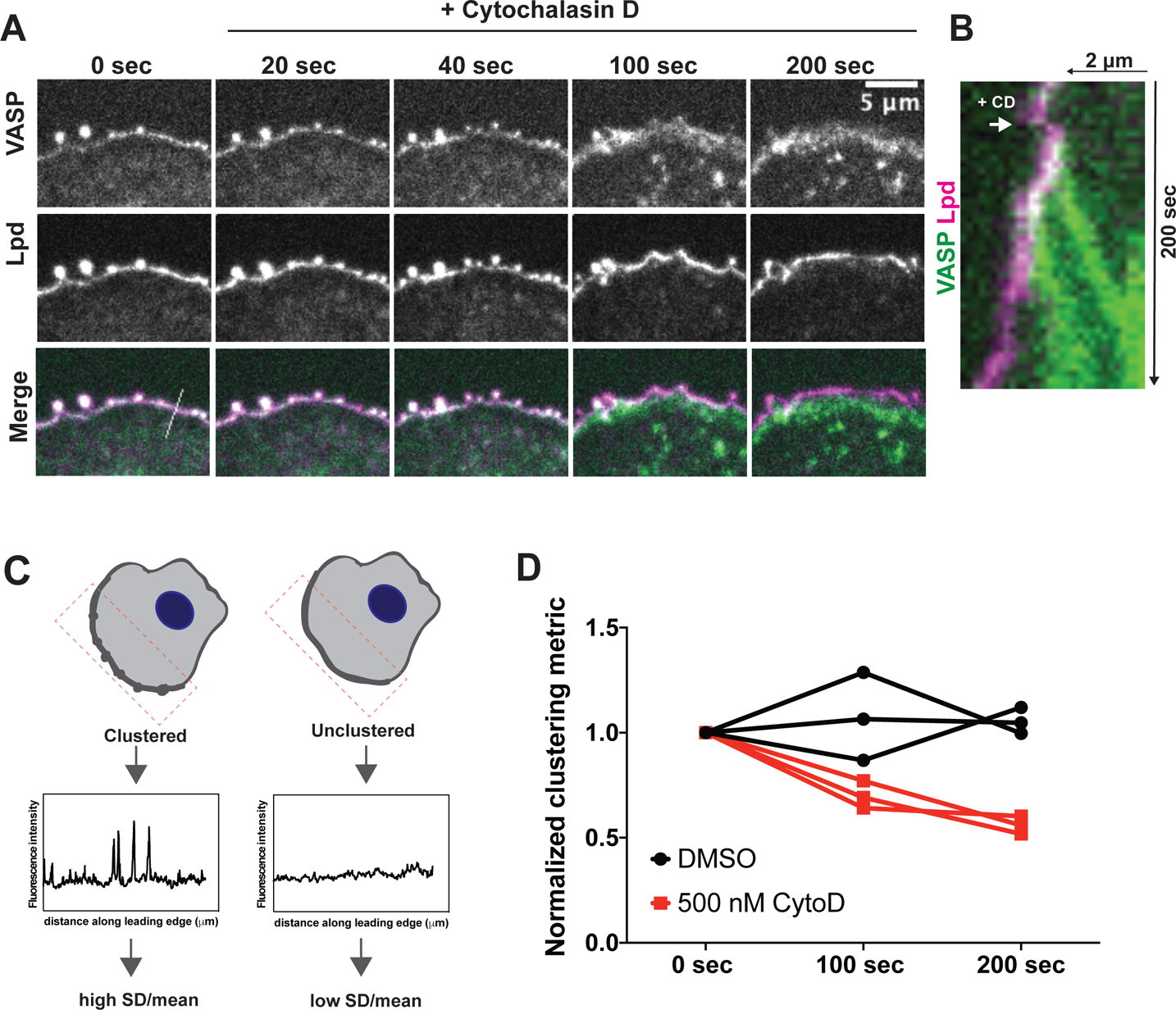
Free barbed ends of actin filaments are required for VASP/Lpd cluster stability. (A) Representative example of a B16F1 cell expressing VASP-eYFP (green) and Lpd-tdTomato (magenta) before and after acute treatment with 500 nM Cytochalasin D. (B) Kymograph of a slice taken perpendicular to the leading edge shows change in membrane localization of VASP (green) and Lpd (magenta) upon addition of CytoD. (C) Schematic of quantification of changes in clustering. The standard deviation along a defined leading edge before (0 sec) and after drug treatment (100 sec, 200 sec) was calculated and normalized to the mean fluorescence intensity. (D) Quantification of clustering of VASP and Lpd at foci following acute treatment with Cytochalasin D or DMSO (vehicle) shows that free barbed ends are required for robust clustering of VASP and Lpd. Clustering is represented by standard deviation along the leading edge ROI before and after treatment. Symbols are averages from three biological replicates, each with >8 cells. Paired t-test of unnormalized data: p=0.02.

## Discussion

The size, shape, and stability of actin-filled pseudopods depends on the dynamic localization of actin regulators in the plasma membrane (Fritz-Laylin et al. 2017; Schmeiser and Winkler 2015; Weiner et al. 2007). Among these regulators is VASP, an actin polymerase that helps construct both sheet-like lamellipodia (Lacayo et al. 2007) and finger-like filopodia (Svitkina et al. 2003; Kwiatkowski et al. 2007; Han et al. 2002). By monitoring the dynamics of VASP and its binding partners in the plasma membrane, we uncovered previously unknown features of the initiation, assembly, and disassembly of filopodia tip complexes. Specifically, we find that VASP clusters assemble at pre-determined sites on the plasma membrane, marked by the presence of its binding partner, lamellipodin. To remain in a cluster, VASP must maintain interactions with both FPPPP-containing ligands *and* free barbed ends of actin filaments. Finally, and most unexpectedly, we find that VASP cluster growth is limited by an intrinsic, size-dependent instability. We propose that the size-dependent instability of VASP clusters reflects the difficulty of coordinating VASP’s interaction with the two binding partners that hold the cluster together: FPPPP-containing proteins and free barbed ends of actin filaments (Figure 7).

**Figure 7.**
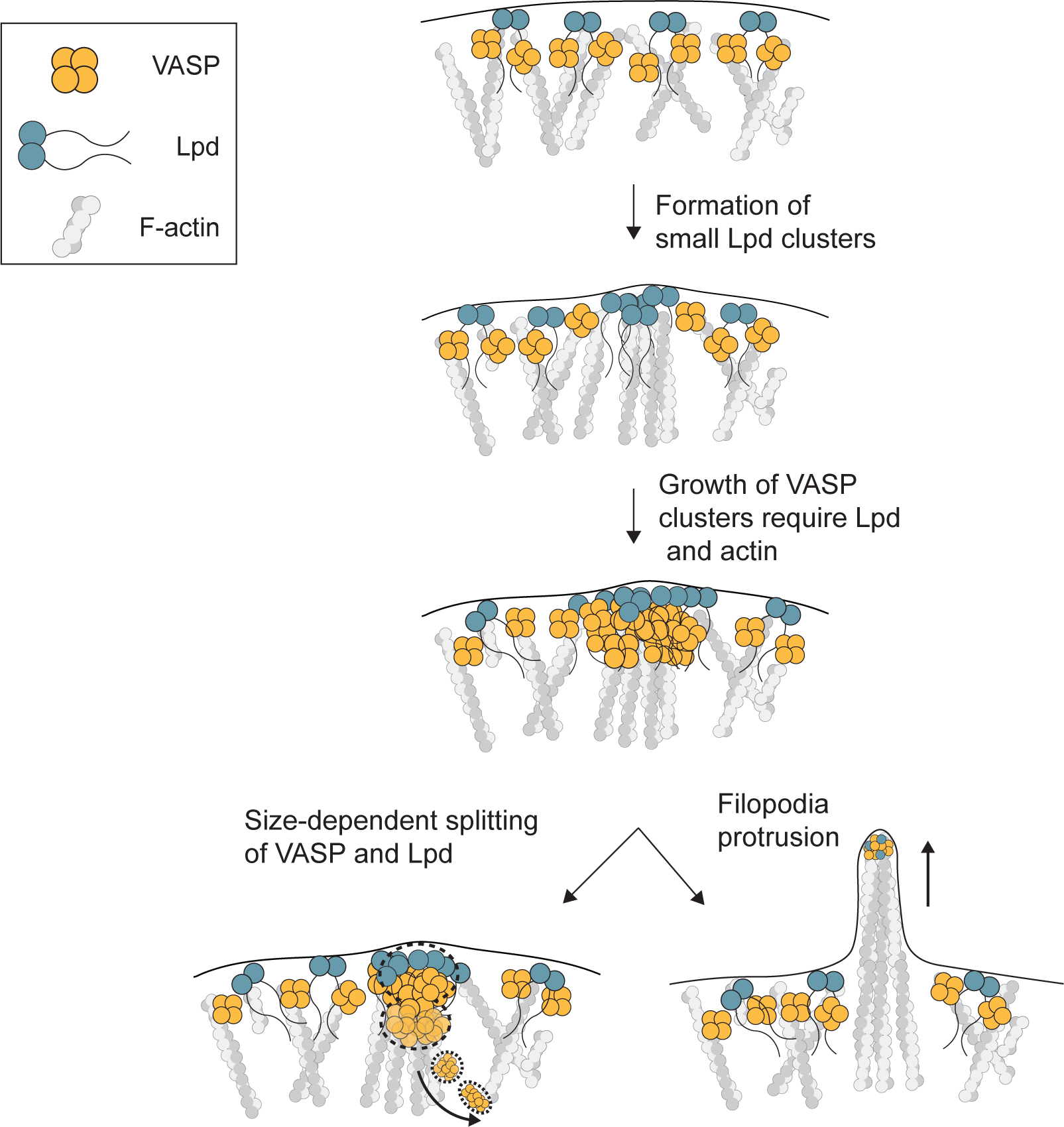
Model. Coalescence of VASP into leading edge clusters arises from small pre-existing lamellipodin clusters. The interaction between VASP and Lpd at clusters are mediated through multivalent interactions between EVH1 domains (VASP) and short F/LPPPP proline motifs (Lpd) that require membrane tethering and dimerization of Lpd. The assembly of VASP clusters involve Lpd and free barbed ends of actin filaments at the leading edge which can protrude into long filopodia or disassemble through size-dependent splitting of VASP into the cytoplasm.

### The role of I-BAR proteins in filopodia assembly

Similar to previous studies, we find that over-expressing the membrane-associated protein, IRSp53 (insulin receptor phosphotyrosine 53 kDa substrate), strongly induces formation of filopodia. Both over-expressed and endogenous IRSp53 localize along the shaft of filopodial protrusions, consistent with the preferential binding of its I-BAR (inverse-Bin-Amphiphysin-Rvs) domain to regions of negative membrane curvature (Yamagishi et al. 2004; Saarikangas et al. 2009). *In vitro*, the I-BAR domain from IRSp53 can tubulate PI(4,5)P_2_-containing vesicles (Mattila et al. 2007; Prévost et al. 2015), and when over-expressed in cells, it can drive formation of tubular membrane protrusions that lack filamentous actin altogether (Yang et al. 2009). Although these results illustrate how easily membrane protrusion can be uncoupled from actin filament assembly, IRSp53 has also been shown to interact, via its SH3 domain, with various actin regulators, including VASP, Mena, N-WASP, mDia1, and Eps8 (Krugmann et al. 2001; Ahmed et al. 2010). Based on its interactions with curved membranes, actin regulators, and signaling molecules such as Cdc42, we and others (Welch and Mullins, 2002; Ahmed et al., 2010) proposed that IRSp53 might trigger filopodium formation by coupling actin assembly to membrane bending. Disanza et al. (2013) further proposed that IRSp53 actually drives formation of filopodial actin bundles by co-clustering with VASP molecules. This attractive idea is generally consistent with the localization IRSp53 to filopodial protrusions, but the strong preference of IRSp53 for curved membranes means it can localize to filopodia and lamellipodia even in Ena/VASP-deficient MV^D7^ cells (Nakagawa et al. 2003). We posed a more stringent test of IRSp53’s involvement in VASP clustering by asking whether IRSp53 associates with nascent VASP foci, or whether it forms forms foci that subsequently recruit VASP. Using fluorescent proteins expressed from the endogenous loci, we found that the coalescence of VASP molecules into foci at the leading edge did not correlate with the presence of IRSp53. Moreover, we observed that VASP foci do not consistently contain detectable amounts of IRSp53 (Figure 2,3). These results fit with recent super-resolution light microscopy studies that found IRSp53 predominantly associated with the lateral edges of filopodia and not consistently enriched at the tip with VASP (Sudhaharan et al. 2019). Our data support the idea that IRSp53 induces protrusion of long filopodia by promoting and stabilizing membrane curvature rather than by clustering VASP tetramers.

### The role of EVH1 ligands in VASP localization and clustering

VASP is recruited to membranes by the interaction of its Ena/VASP Homology 1 (EVH1) domain with proteins that contain F/LPPPP motifs (Bear et al. 2002). This was most clearly demonstrated by Bear et al. (2002), who found that VASP can be displaced from leading edge membranes by removing the EVH1 domain or ectopically expressing F/LPPPP-containing proteins on intracellular membranes. In addition to lamellipodin, several other proteins with EVH1-binding motifs are found on leading edge membranes. These include Mig10 and RIAM which, together with lamellipodin, define the MRL protein family (Coló et al. 2012).

When we compared the dynamics of endogenous VASP and lamellipodin during filopodia initiation, we found that VASP clusters form on pre-existing foci of lamellipodin. The small lamellipodin puncta form independently of VASP, but ‘ripen’ and grow much larger when they begin to accumulate VASP. This observation reveals that small lamellipodin puncta are precursors to larger VASP/lamellipodin clusters that concentrate at the tips of nascent filopodia bundles and raises the question of how small lamellipodin puncta form in the first place. Two possibilities are: (i) lamellipodin can independently form small homo-oligomers, which is supported by a structural study that identified two dimerization motifs at its N-terminal RAPH domain (Chang et al. 2013) and (ii) lamellipodin foci mark the position of larger complexes that contain other, as yet unidentified proteins.

Our results provide compelling evidence that VASP/Lpd interactions help hold filopodial tip complexes together. We found, for example, that monomeric, C-terminal Lpd mutants disrupt formation of VASP clusters. We also identified the key features of Lpd that promote VASP clustering: dimerization and EVH1-binding. Surprisingly, the N-terminal Ras-association (RA) plekstrin homology (PH) signaling domain of Lpd appears largely dispensable for VASP clustering as we could functionally replace it with a Lyn_11_ membrane tether and a leucine zipper dimerization domain. Based on our results we propose that the primary function of the RAPH domain is membrane localization rather than cluster formation. This proposal agrees with several studies demonstrating that the PH domain Lpd recruits the protein to the plasma membrane by binding PI(3,4)P_2_ (Bae 2010, Venkatareddy 2011). Our proposal also fits with a recent structural study of Lpd’s N-terminal region which suggests that its dimeric RAPH domain has lower affinity for Ras GTPases than individual RA domains found in other proteins, and may make minor contributions to clustering on membranes (Chang et al. 2013).

While multiple lines of evidence indicate that FPPPP-EVH1 interactions are responsible for VASP localization to the leading edge, Dimchev et al. reported that knocking out lamellipodin by CRISPR-Cas9 in B16F1 cells does not significantly change VASP recruitment to lamellipodia (Dimchev et al. 2019; Bear et al. 2002). This result suggests the existence of redundancy or compensation in the expression of FPPPP-containing proteins. For example, Dimchev et al (2019) also reported that leading-edge localization of a member of the WAVE Regulatory Complex, Abi1, which binds EVH1 domains through an alternate proline motif was increased upon Lpd knockout and may allow it co-cluster with VASP in the absence of lamellipodin (Chen et al. 2014; Dimchev et al. 2019). In addition, lamellipodin is just one member of the MRL family of proteins, which all interact with VASP through FPPPP proline motifs and could potentially replace Lpd’s VASP-specific functions at the leading edge.

### Size control of VASP/Lpd clusters

To maintain a dynamic and dispersed distribution along the leading edge, VASP/Lpd clusters continuously undergo assembly and disassembly. Growth of clusters through gradual accumulation of monomers or abrupt fusion events must be counterbalanced by a disassembly mechanism to prevent the total collapse of VASP into a few large, static foci at the leading edge. By analyzing VASP cluster size fluctuations, we discovered a novel disassembly mechanism whereby VASP/Lpd clusters undergo size-dependent splitting events. During splitting, a bolus of VASP detaches from a cluster and moves into the cytoplasm while an equimolar amount of Lpd redistributes evenly back into the plasma membrane. The larger the VASP/Lpd cluster grows, the more unstable it becomes and the instability grows as the approximate square of the the cluster size. The fact that these splitting events occur perpendicular to the plasma membrane while fusion events occur parallel to the membrane indicates that splitting is not simply the reverse of fusion, but a separate process with a different underlying molecular mechanism. In addition to maintaining dynamic VASP clusters, splitting might help relocalize VASP tetramers to other parts of the cell. For example, in some cases where nascent focal adhesions are proximal to the leading edge, we observed blobs of VASP splitting from the a plasma membrane focus and almost immediately becoming immobilized in the elongated shape of a focal adhesion. As several focal adhesion proteins (such as vinculin and zyxin) directly bind VASP through the F/LPPPP-EVH1 interaction module, it is possible that fission of large VASP/Lpd clusters helps deposit VASP directly onto nascent focal adhesions.

## Materials and methods

### Constructs and reagents

The following primary antibodies and dyes were used for staining: polyclonal rabbit antibody against IRSp53 (4 μg/ml, Atlas Prestige antibodies, HPA023310), monoclonal rabbit antibody against VASP (1:100 dilution, Cell Signaling, #3132), Alexa-647 phalloidin (Thermo Fisher #A22287). Various Lpd constructs were derived from an original plasmid encoding human Lpd^1-1250^provided by Matthias Krause (King’s College, London) and subsequently cloned into a pCMV-SNAP vector.

### Cell culture

B16F1 (ATCC-CRL-6323) mouse melanoma cells were cultured in Dulbecco’s modified Eagle medium (DMEM) with 4.5 g/mL glucose and supplemented with 10% fetal bovine serum (FBS, Gibco certified, US) and penicillin/streptomycin. Cells were maintained at 37°C and 5% CO_2_ and split every 2-3 days.

Lipid-based transient transfection was performed using Lipofectamine 3000. For the 6-well plate format, 7.5 μl Lipofectamine 3000 was combined with 1 μg DNA and 5 μl P3000 reagent. The transfection mixture was incubated with 50-70% confluent cells for 8-16 hours and then replaced with fresh media. Cells were assayed or harvested 48 hours later.

### Cell line generation

The VASP locus was tagged with eYFP at the C terminus using CRISPR/Cas9-mediated genome engineering in B16F1 cells. The oligonucleotide guide sequences used to cut the targeted genomic sites are listed in Table 1 below. Cas9 and single-guide RNA were expressed using pX330 as previously described (Ran et al. 2013). Both pX330 and the eYFP-neomycin donor plasmid were a gift of Kara McKinley (UCSF). pX330 and the donor plasmid were co-transfected into B16F1 cells at 1.25 μg each and selected after 72 hours with 0.8 mg/ml G418 for 2 weeks. Monoclonal cell lines were generated by fluorescence activated cell sorting (FACS) and verified by sequencing and microscopy. Lpd was sequentially tagged with tdTomato-puromycin at the C terminus using the same strategy as above. eYFP-VASP and tdTomato-Lpd double knock-in cell lines were selected using 2 ng/μl puromycin for 5 days and subsequently sorted for monoclonal colonies by FACS.

**Table 1:**
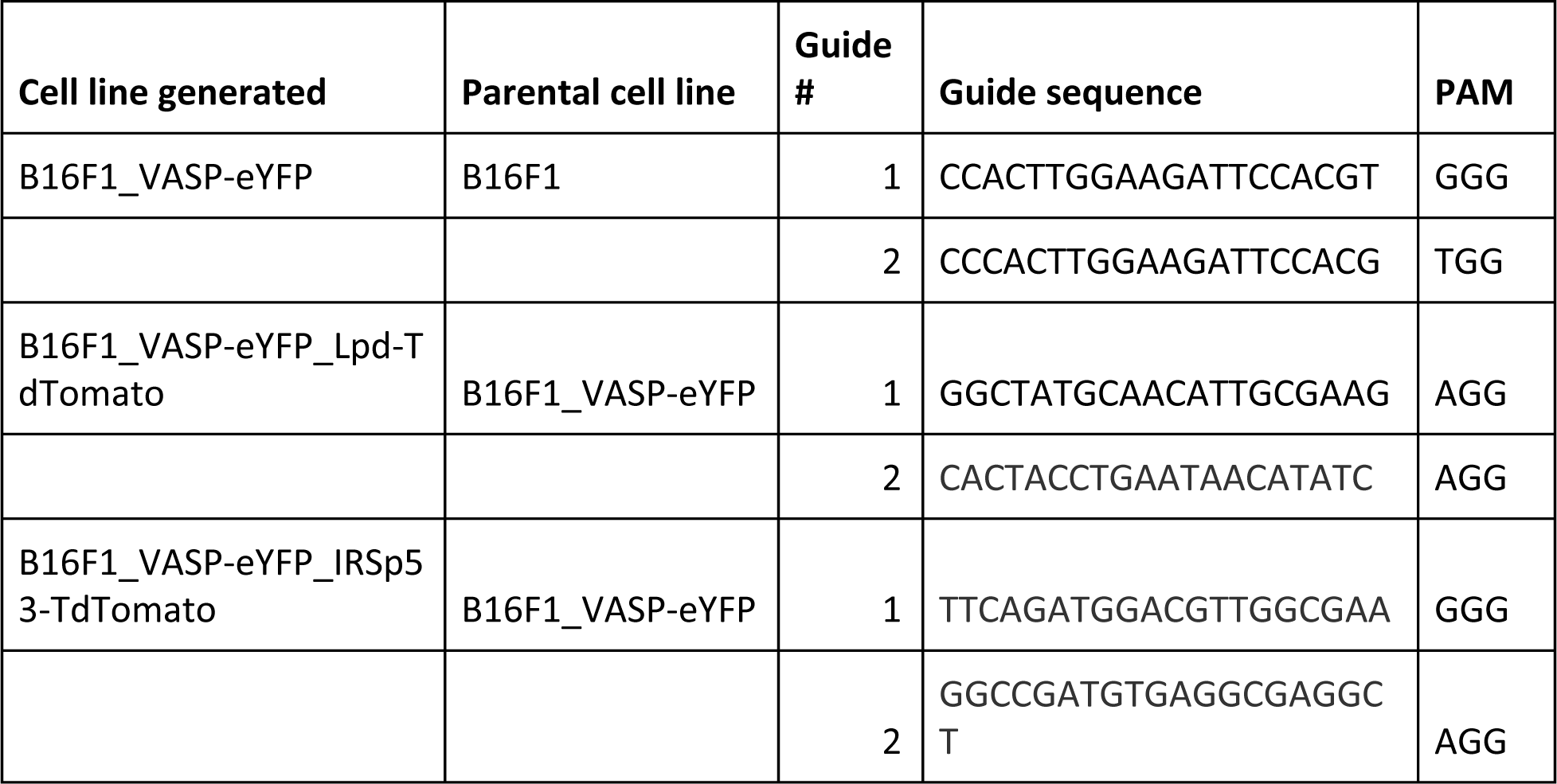
Knock-in guide sequences

### Sample preparation for microscopy

Glass chambers of varying sizes were cleaned by plasma cleaning for 5 minutes, coated with 50 μg/ml mouse laminin (Sigma #L2020) diluted in PBS, and incubated overnight at 4°C. The day of imaging, laminin-coated glass was rinsed with PBS and replaced with imaging media (Ham’s/F12 media + 10% FBS + 50 mM HEPES). B16F1 cells were trypsinized and plated on laminin-coated glass and began to spread within 30 minutes. Cells transfected with SNAP-fusion proteins were labeled with 50 nM Janelia Fluor SNAP_646_-ligand diluted in imaging media for 1 hour upon plating on laminin-coated glass. Janelia Fluor SNAP ligands were a generous gift of Luke Lavis (Janelia Farms). Prior to imaging, the cells were washed with PBS and replaced with fresh imaging media.

### Microscopy

Cells were imaged by confocal spinning disk microscopy using a 60x Nikon Plan Apo TIRF (NA 1.49) objective on a Nikon Eclipse microscope fitted with an Andor iXon emCCD camera and controlled by Micromanager 1.4. Cells were maintained at 37°C and 5% CO_2_ during imaging using an Okolab stage top incubator and controller. Image processing was performed using ImageJ/Fiji and custom Matlab code.

For experiments in which the barbed ends of actin filaments were blocked using Cytochalasin D (Sigma #C8273), B16F1 cells were first plated on laminin-coated glass as described above. To obtain a baseline before drug addition, we imaged cells for 30 seconds then added 500 nM Cytochalasin D (or DMSO) directly on the microscope stage for acute perturbation. The drug was prepared at 2x concentration in imaging media before addition to cells to achieve a final concentration of 500 nM.

### Image analysis

To track individual VASP/Lpd clusters, we developed custom Matlab code to identify fluorescent protein clusters through particle tracking in a time-lapse image stack. All movies analyzed contained 40 frames and were acquired at a frame rate of 15 frames/minute. Image acquisition parameters were identical across all experiments. For each frame, the function creates an ROI around the spot and calculates the background-subtracted, integrated intensity of a 2D Gaussian spot centered on the point. We refer to this metric as “time-averaged cluster intensity” in the text and figures. The intensity fluctuations of both VASP and lamellipodin were represented by the standard deviation in integrated cluster intensity over the fixed imaging time window.

To generate adaptive kymographs to follow fluorescent protein dynamics along a specified leading edge, we developed a Matlab function that takes in two color time-lapse data sets, uses the marker fluorescence of a designated fluorescence channel to identify the cell edge and, with user input, creates a dynamic region of interest around the cell edge. Tracking and plotting parameters include the following: gaussian blurring used to smoothen images; threshold to find the cell edge; absolute maximum intensity of channel 1 for plot display; absolute maximum intensity of channel 2 for plot display. The moving leading edge is aligned frame-by-frame using autocorrelation. Using this region of interest, the function creates a kymograph for the original channel (VASP) and a second, independent channel (i.e. Lamellipodin or IRSp53).

### Clustering quantification

VASP clustering at the leading edge was quantified by measuring the background-subtracted fluorescence intensity over an 8-point line width ROI traced along the leading edge in ImageJ/Fiji. The clustering metric was calculated as the standard deviation along the leading edge ROI normalized to the average fluorescence intensity (SD/mean). For paired samples (such as in the cytochalasin D treatments), the drug treated condition was normalized to the control condition.

## Acknowledgements

We thank Kara McKinley, Scott Hansen, Luke Lavis, and Samuel Lord, and other members of the Mullins lab for reagents, experimental advice, and feedback on the manuscript. We thank Ron Vale, Sophie Dumont, Geeta Narlikar, Dan Fletcher, and Orion Weiner for helpful conversations. This work was supported by the National Institute of General Medical Sciences of the National Institutes of Health (R35-GM118119 to R.D. Mullins), by the Howard Hughes Medical Institute Investigator program (R.D. Mullins), and by a predoctoral fellowship from the American Heart Association (17PRE33630125 to K.W. Cheng).

## Supplemental videos legend

**Video 1: Endogenous VASP-eYFP localization and dynamics at the leading edge.**

B16F1 cells expressing VASP-eYFP from the endogenous gene locus imaged by spinning disc confocal microscopy at 4-second time points.

**Video 2: VASP-eYFP foci coalesce at the tips of filopodia actin bundles.**

B16F1 VASP-eYFP (green) cell line with over-expressed SNAP_JF646_-actin (magenta) imaged by spinning disc confocal at 4-second time points.

**Video 3: Lamellipodin localizes to dynamic VASP clusters and displays coupled dynamics at the leading edge.**

B16F1 VASP-eYFP (green) and lamellipodin-tdTomato (magenta) cell line imaged by spinning disc confocal at 4-second time points.

**Video 4: IRSp53 displays punctate membrane localization but does not concentrate in VASP foci.**

B16F1 VASP-eYFP (green) and IRSp53-tdTomato (magenta) cell line imaged by spinning disc confocal at 4-second time points.

**Video 5: Size-dependent splitting of VASP from VASP/lamellipodin clusters at the leading edge.**

B16F1 VASP-eYFP (green) and lamellipodin (magenta) cell line imaged by spinning disc confocal at 4-second time points. White arrow (left) indicates the parental cluster within the leading edge membrane; white arrow (right) indicates the VASP that splits away towards the cytoplasm.

**Video 6: Capping barbed ends disrupts clustering of VASP and lamellipodin.**

B16F1 VASP-eYFP (green) and lamellipodin-tdTomato (magenta) cell line were treated with 500 nM Cytochalasin D at a 30-second time point and imaged by spinning disc confocal at 4-second time points.

## Supplemental Figure Legends

**Supplementary Figure 1.**
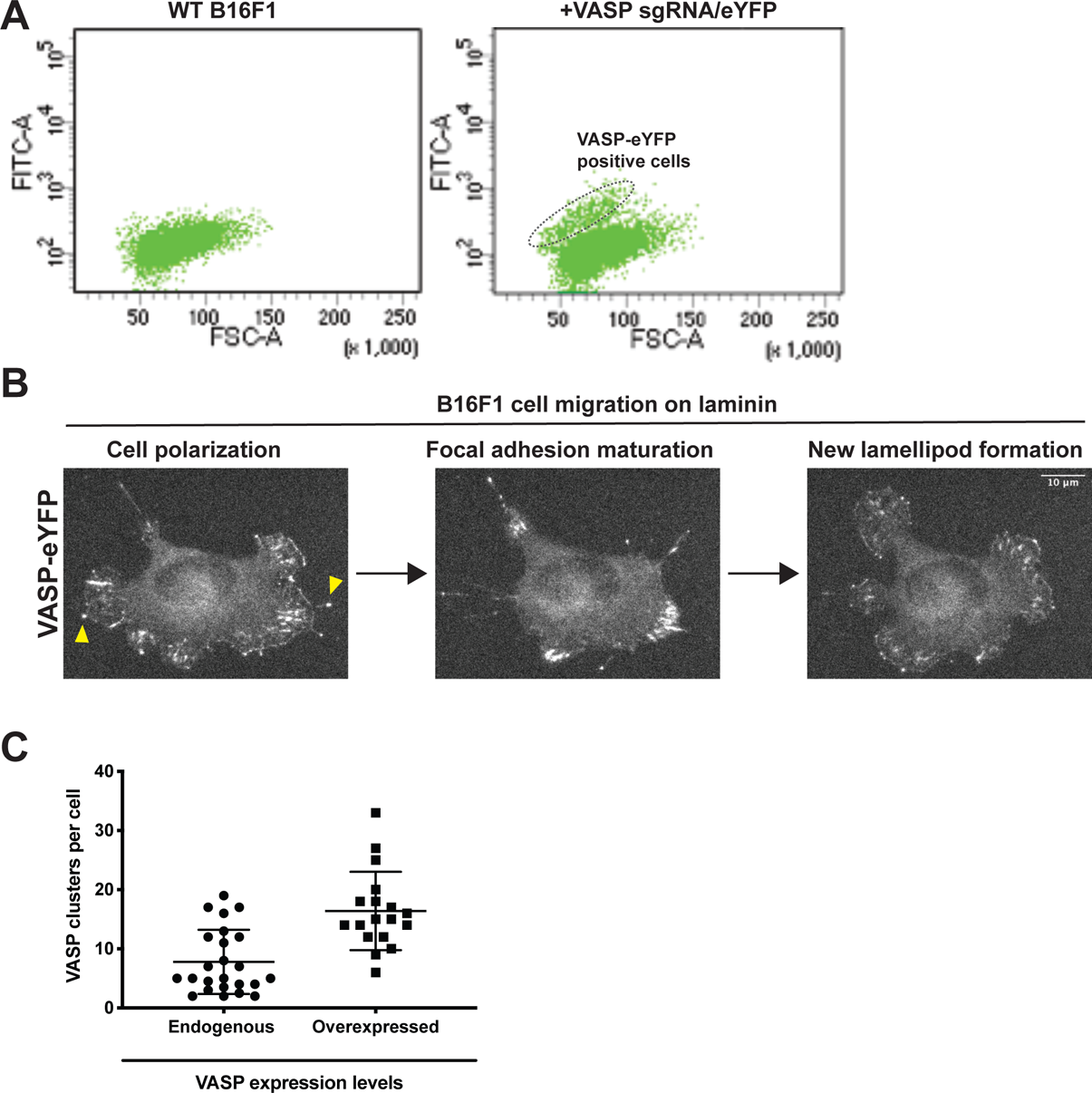
Generation of monoclonal VASP-eYFP B16F1 mouse melanoma cell line. (A) Dot plot from fluorescence activated cell sorting (FACS) of B16F1 cells transfected with Cas9/VASP-specific sgRNA and C-terminal eYFP tag repair template compared to WT B16F1 cells. (B) Endogenously tagged VASP-eYFP B16F1 cells display normal migratory behavior on laminin coated glass. Cell polarization leads to formation of a leading edge with correct VASP localization at lamellipodia, filopodia, and focal adhesions (Yellow arrowheads). Over time, the cell creates mature focal adhesions which is followed by new lamellipod formation and migration in a new direction. (C) The number of VASP clusters per cell in endogenously tagged VASP-eYFP cells or overexpressed GFP-VASP were calculated by counting the number of VASP clusters per 160 second time window using a Z-projection of the movie stack for easy visualization.

**Supplementary Figure 2.**
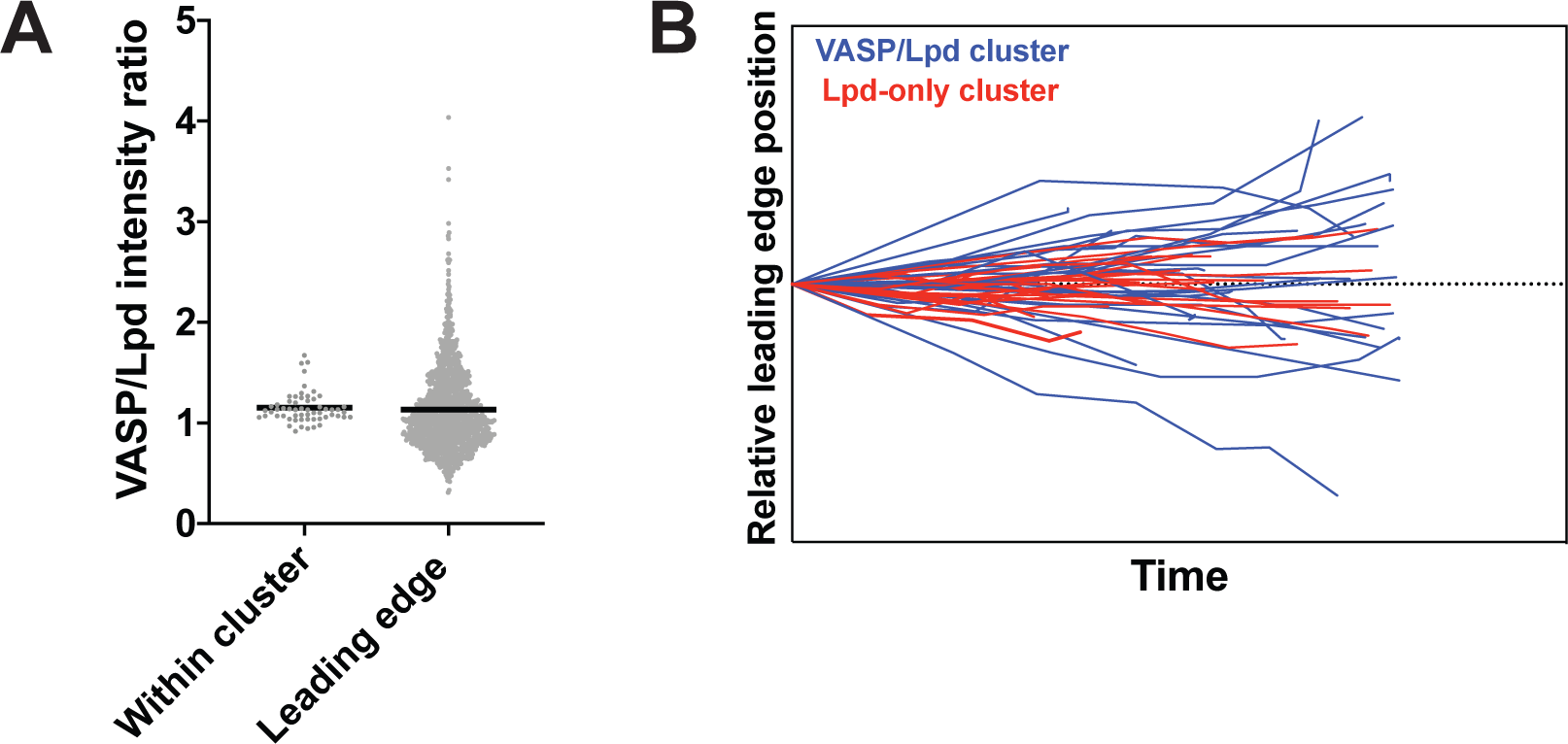
Distinct dynamics within VASP/lamellipodin clusters compared to the entire leading-edge. (A) The ratio of VASP to lamellipodin intensities within clusters were and compared to the ratio of VASP to lamellipodin intensities along every point on the leading edge of B16F1 cells. (B) Quantification of lateral skating movement of VASP/Lpd clusters (30 cluster tracks) vs. clusters that only contain Lpd (23 cluster tracks) along the leading edge. The starting position was normalized to begin at the same point for all tracks.

**Supplementary Figure 3.**
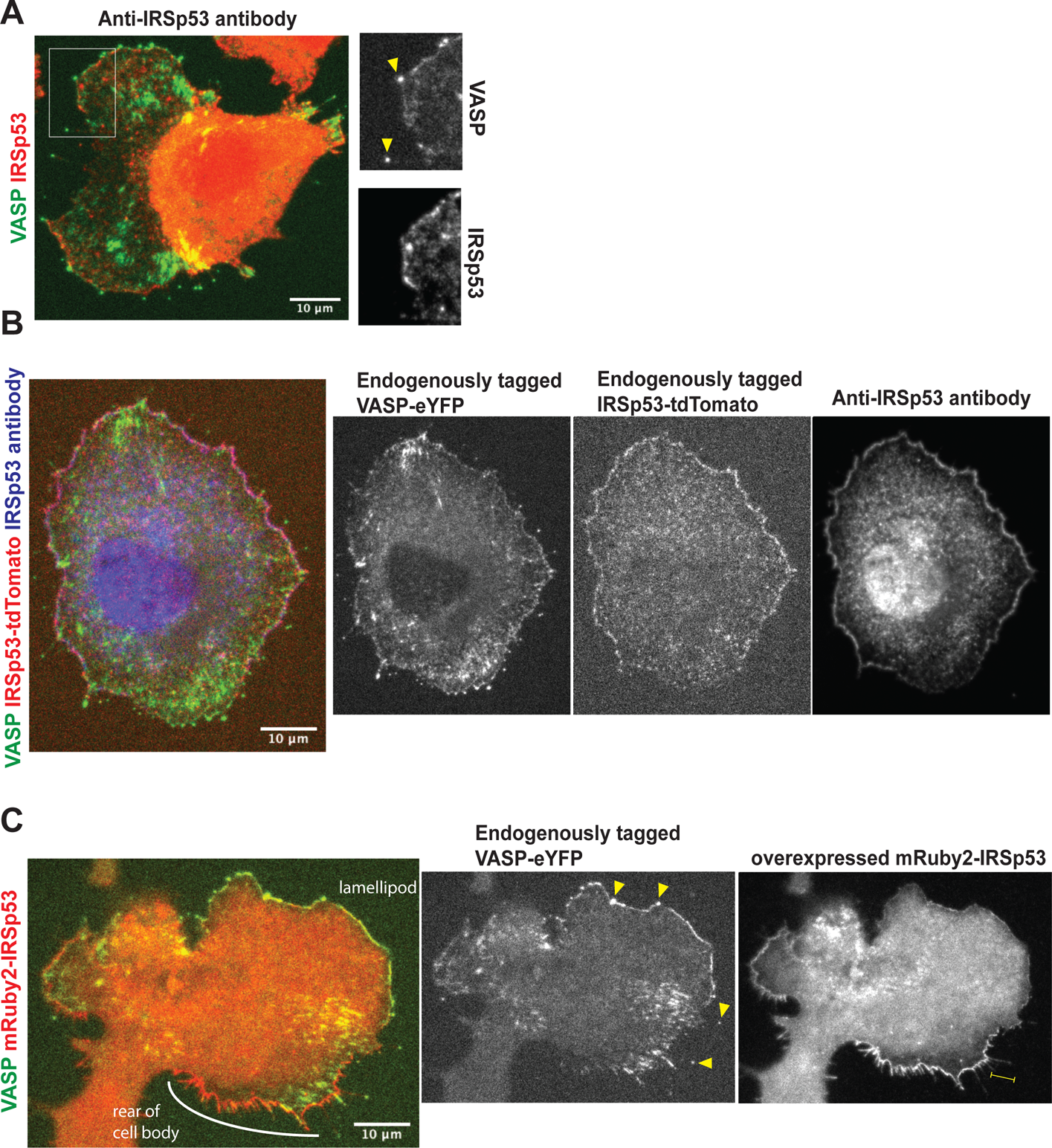
Localization of endogenous and over-expressed IRSp53. (A) Immunofluoresence of endogenous IRSp53 (red) in VASP-eYFP (green) tagged B16F1 cells. Arrowheads: VASP-eYFP foci; yellow bar: filopodia shaft. (B) Localization of overexpressed, N-terminally tagged mRuby2-IRSp53 in the context of the VASP-eYFP B16F1 CRISPR knock-in cell line. Overexpressed mRuby2-IRSp53 concentrated at the plasma membrane and in the shaft of negative curvature tubules such as filopodia and retraction fibers but not leading edge VASP clusters. (C) Verification of double CRISPR knock-in VASP-eYFP/IRSp53-tdTomato B16F1 cells by immunostaining with a polyclonal IRSp53 antibody (Atlas Prestige from Sigma Aldrich) and secondary Alexa 647 antibody (Life Technologies).

**Supplementary Figure 4.**
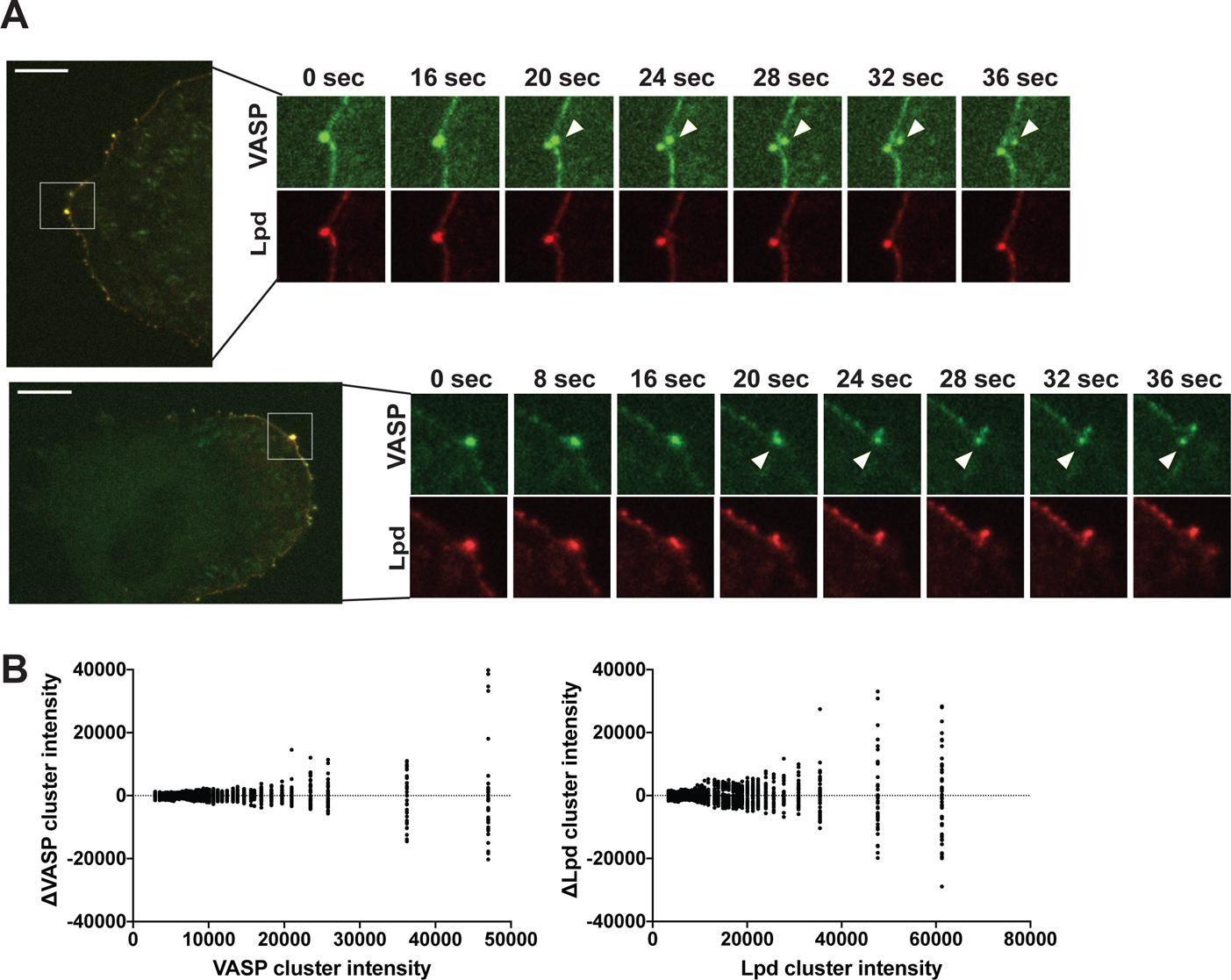
Further characterization of size-dependent splitting of VASP/Lpd clusters at the leading edge. (A) Two additional examples of VASP/Lpd cluster splitting events. Insets represent frames capturing the splitting event with white arrowhead tracking the chunk of VASP that is left behind in the cytoplasm upon splitting. (B) Alternative visualization strategy of the size-dependency of VASP/Lpd cluster instability. The X axis represents the time-averaged cluster size over a 160 second time window given by the integrated fluorescence intensity of either VASP-eYFP (left subplot) or Lpd-tdTomato (right subplot). The Y axis represents the relative deviations of the cluster size at every time point over the 160 second imaging window (Δ cluster size = time-averaged cluster size - cluster size(t)).

